# Exercise intensity modulates the human plasma secretome and interorgan communication

**DOI:** 10.1101/2025.10.22.683699

**Authors:** Luke Olsen, Javier Botella, Douglas Barrows, Ethan Romero, Kaitlyn Baird, Mutsumi Katayama, Ece Kilic, Christopher Peralta, Henry Sanford, Laurie Farrell, Christopher L. Axelrod, Kaja Plucińska, Jeanne Walker, Lu Yan, Katie Fredrickson, Olivier Pourquie, Jeremy M. Robbins, Ekaterina V. Vinogradova, John P. Kirwan, Juleen R. Zierath, Anna Krook, Robert E. Gerszten, David J. Bishop, Paul Cohen

## Abstract

Exercise is recognized as first-line therapy for many cardiometabolic diseases, including obesity, type 2 diabetes, and hypertension. Despite the abundant health-promoting effects of exercise, in-depth characterization of circulatory factors that mediate these benefits in humans remains incomplete. Moreover, how different modes and intensities of exercise uniquely regulate these processes is unclear. Here, we address these questions by conducting a multi-cohort human exercise intervention, incorporating sprint-interval exercise (SIE) and moderate-intensity exercise (MIE) to analyze intensity-dependent regulation of interorgan crosstalk. We find that exercise intensity distinctly influences the plasma proteome and metabolome in both untrained and trained participants. SIE led to immediate and robust changes to the plasma proteome, whereas MIE resulted in delayed secretory kinetics. By leveraging large, multi-organ gene and protein expression datasets, in combination with *in vitro* and *in vivo* tissue sampling, we map the differentially regulated proteins to their predicted tissue of origin and destination. We find that adipocytes are particularly sensitive to exercise intensity, undergoing broad transcriptomic remodeling following *in vitro* incubation with SIE as compared to MIE plasma. These findings underscore the integrated whole-body response following acute exercise and highlight exercise intensity as a key factor influencing interorgan communication.

## Introduction

The metabolic benefits of regular exercise reflect integrated multi-organ coordination, spanning distant tissues and diverse cell-types. Secreted metabolites, lipids, peptides, and proteins govern exercise-regulated interorgan crosstalk and contribute substantially to exercise-dependent adaptations^1^. In addition to upregulating secretory factors, termed exerkines^2^, exercise also downregulates the levels of deleterious factors. Collectively, these circulating mediators (the secretome) regulate multiple aspects of metabolism, such as promoting adipose tissue remodeling by reducing fibrosis while increasing vascularization and innervation^3,4,5^, enhancing whole-body glucose handling through metabolic rewiring in skeletal muscle^6^, and improving cognition through stimulating neurogenesis and inhibiting neuroinflammation^7^.

Increasing evidence suggests that not all exercise is equal. Distinct exercise intensities, such as sprint-interval exercise (SIE) or moderate-intensity exercise (MIE), elicit intensity-dependent, body-wide adaptations^8^, that may in part be driven by differences in secretory profiles^9,10^. Supporting this, N-lactoyl-phenylalanine (Lac-Phe), a recently described obesity-mitigating metabolite, is secreted following exercise in an intensity-dependent manner, with greater secretion following sprint exercise compared to resistance or endurance exercise^11,12^. Similarly, many soluble proteins, such as interleukin-6 (IL6) and growth differentiation factor 15 (GDF15), are also secreted in an intensity-dependent manner, resulting in broad and context-dependent metabolic changes^13,14,15^. Characterizing intensity-dependent secretory responses may uncover how sprint-interval exercise — often lasting less than five minutes in total work — results in similar, if not superior, whole-body metabolic adaptations compared to longer duration low-or moderate-intensity exercise^16^.

Despite the growing evidence suggesting intensity-dependent regulation of the plasma proteome and metabolome, in-depth comparisons following acute and chronic training interventions remain limited. Additionally, the identity of the tissue of origin and destination following different styles of exercise remains largely unexplored in humans. To address these knowledge gaps, we characterized the plasma proteome and metabolome following a single session of SIE and MIE in participants before and after 8 weeks of training (hereafter termed untrained and trained). Incorporating a multiplexed, antibody-based protein detection platform, in combination with untargeted metabolomics, we uncover striking intensity-dependent changes to the plasma secretome — observed in both untrained and trained participants — and further characterize their predicted tissue of origin and target tissue. These findings provide new insight into diverse interorgan communication streams following exercise in humans and highlight the metabolic benefits, and potential limitations, of incorporating diverse exercise intensities into a regular training program.

## Results

### The plasma proteome is differentially regulated by exercise intensity

To characterize acute changes to the plasma proteome following two distinct types of exercise, we collected plasma before (rest), immediately following (0h), and 3 hours following (3h) moderate-intensity exercise (MIE; N=9) or sprint-interval exercise (SIE; N=10) in young, active, metabolically healthy males^17^. MIE consisted of 90-minutes of continuous cycling at 90 to 100% of the predetermined first lactate threshold, while SIE consisted of 6 sets of 30-second all-out cycling at a resistance of 0.075 kg•kg^-1^ body mass, with 4-minute rests between sets (Figure 1A). As anticipated and by design, blood lactate levels and pH were increased and decreased, respectively, immediately following SIE, with minimal change following MIE^17^.

**Figure 1.**
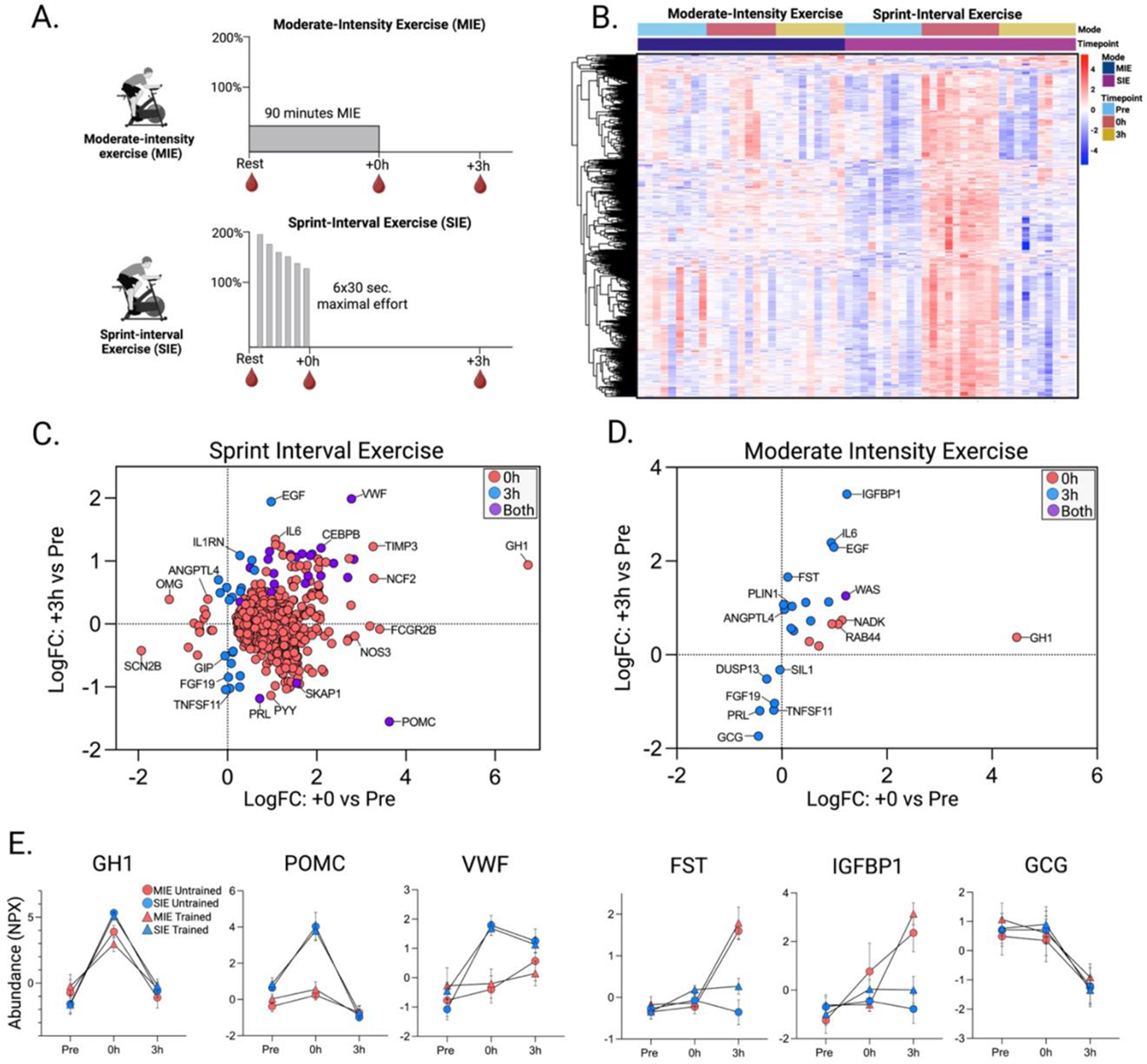
Acute SIE and MIE uniquely regulate the plasma proteome. **A**. Experimental design for moderate-intensity exercise (MIE) and sprint-interval exercise (SIE) detailing exercise duration and sample collection (blood) timepoints. **B**. A heatmap showing z-scores of mean Olink NPX values for all differentially regulated proteins between Pre and 0h (immediately post) or 3h (3-hours post) following MIE or SIE. Concordance analysis of the log fold change (LogFC) for differentially regulated proteins between Pre vs. 0h and Pre vs. 3h for SIE (**C.**) and MIE (**D.**). **E**. Plasma protein abundance of representative proteins in untrained and trained participants following either SIE or MIE. Values are raw Normalized Protein values (NPX). Only participants with values in untrained and trained conditions are displayed (N=6-7 per condition). Error bars indicate standard error of the mean.

To efficiently analyze the plasma proteome across its broad dynamic range, we used the antibody-based detection technology Olink Explore^18^, which detected and quantified 2,884 proteins across all participants and timepoints. Critically, this pipeline reliably detected proteins within the ng/mL to pg/mL range, including interleukin-1ß (IL-1ß), interleukin-16 (IL-16), and IL-6. Principal Component Analysis (PC) revealed intensity-dependent changes to the plasma proteome, with samples collected immediately following SIE shifting along both PC1 and PC2, and returning to resting levels 3h post. In contrast, MIE showed no time-dependent clustering regardless of timepoint (Figure S1A and S1B).

SIE robustly altered the plasma proteome, with nearly a quarter (714 proteins) of the total detected proteins changing immediately following exercise (FDR<0.05; Figure 1B and 1C). Of these, >98% were increased in abundance relative to resting levels, including known exercise-responsive proteins growth hormone 1 (GH1), tissue inhibitor of metalloproteinase 3 (TIMP3), von Willebrand factor (VWF), and fatty acid binding protein 5 (FABP5). The number of differentially regulated proteins decreased nearly 20-fold 3h post, highlighting rapid changes to the plasma proteome following SIE. In stark contrast, only 7 proteins were differentially regulated immediately following MIE, which increased to 19 proteins 3h post (Figure 1D). MIE-regulated plasma proteins included follistatin (FST), insulin-like growth factor binding protein 1 (IGFBP1), and perilipin 1 (PLIN1). Despite minimal overlap between exercise modes, both interventions increased GH1 and NAD-kinase (NADK) 0h post, while decreasing prolactin (PRL), receptor activator of nuclear factor-ß ligand (RANKL), and fibroblast growth factor 19 (FGF19) 3h post (Table S1).

To determine whether the observed changes to the plasma proteome are specific to the untrained state or persist in trained subjects, we conducted an 8-week training regimen with a subset of the original participants, incorporating their respective mode of exercise: sprint-interval training (SIT; n=7) and moderate-intensity training (MIT; n=6). Following 8 weeks, we conducted an identical acute exercise test and plasma processing pipeline as described above^17^. Intriguingly, plasma protein dynamics displayed similar temporal responses following acute exercise in the trained state, with an immediate and robust increase following SIE, and a delayed increase following MIE (Figure S2A–S2C) — findings supported by significant correlation between untrained and trained datasets (Figure S2D–S2G). Moreover, many of the shared and intensity-dependent changes persisted in the trained state, including GH1, VWF, IGFBP1, FST, POMC, and GCG (Figure 1E), highlighting the quantitative and temporal intensity-dependent changes to the plasma proteome following a single session of exercise in both untrained and trained participants.

We next assessed the generalizability of our findings to commonly prescribed treadmill running, specifically to determine whether the low number of differentially regulated proteins following MIE is a result of exercise intensity, exercise modality (e.g., cycling), or a combination of the two. To test this, we collected and analyzed plasma samples from 11 healthy, active males before and immediately following 2 hours of running at 60% of their VO^2max19^ — an exercise regimen similar to MIE cycling. Using an identical protein processing pipeline as described above, we detected a total of 2,787 proteins, 111 of which were differentially regulated following exercise. Of these, nearly all (109 proteins) increased relative to baseline (Figure S2H). Most of the shared proteins immediately following exercise were between SIE cycling and running, including ANGPTL4, VWF, FABP4, and ADAMTS1, suggesting moderate-intensity running perturbs the plasma proteome to a greater extent than moderate-intensity cycling, at least at the 0h timepoint (Figure S2I and S2J). Nonetheless, the number of differentially regulated proteins following moderate intensity running remained markedly lower than following SIE, further highlighting the robust and dynamic remodeling of the plasma proteome following sprint-interval exercise compared to running or cycling at moderate intensities.

Finally, we tested whether any differentially regulated plasma proteins were associated with future health or disease outcomes. To do this, we leveraged a UK Biobank plasma proteome-phenome resource comprising 53,026 human participants and 1,066 incident- and prevalent- based diseases^20^. Across the 741 exercise-regulated proteins, we identified 46,993 protein-disease associations (Bonferroni-adjusted p-value <0.05). As expected, >95% of these associations had hazard or odds ratios (HR/OR) greater than 1, consistent with prior work showing acute exercise stimulates a transient inflammatory state^21^. To identify proteins potentially protective against metabolic disease, we filtered the 46,993 associations for those with HR/OR <1 and then restricted this list to phenotypes of interest: “metabolic disorders”, “obesity”, and “type 2 diabetes”. In total, 33 proteins remained (Figure S3; Table S2). Almost all – 32 out of 33 proteins – were differentially regulated following SIE at either 0h (28 proteins) or 3h (4 proteins). In contrast, MIE regulated three of the proteins: TNFSF11, FGF19, and FST. Notably, ADGRG2, FGFBP1, and MXRA8 — all increasing immediately following SIE — consistently showed low HR/OR across metabolic states. In fact, in a UK Biobank subgroup (n=32,549), plasma levels of ADGRG2 and MXRA8 were positively associated with regular physical activity^22^, suggesting their potential benefit extends beyond the acute exercise window.

### Intensity-dependent regulation of the plasma metabolome following SIE or MIE

While proteins have classically been studied for their endocrine function, secreted metabolites have become increasingly appreciated for their potential to act as molecular transducers, as highlighted by the characterization of exercise-responsive metabolites 2-hydroxybutyrate, Lac-Phe, ß-aminoisobutyric acid, and 12,13-diHOME^11,23,24,25^. To identify additional metabolites secreted in an intensity-dependent manner, we performed untargeted polar plasma metabolomics following SIE or MIE, and characterized the differential regulation of both annotated and unannotated metabolites (see methods).

In contrast to the plasma proteome, principal component analysis revealed intensity-dependent and time-dependent clustering of the plasma metabolome following MIE and SIE (Figure 2A). In total, SIE resulted in the differential regulation of 203 metabolites immediately following exercise, including well-described exercise-responsive metabolites lactate, succinate, malate, and pyruvate (Figure 2B). We also found lactate-derived metabolites increased after SIE, most notably Lac-Phe, which displayed similar temporal dynamics to circulating lactate (Figure 2B). Moreover, kynurenic acid — a metabolite secreted by skeletal muscle which signals to adipose tissue^26^ — increased following SIE, corresponding with a rise in circulating glycerol. At 3h following SIE, 199 metabolites were differentially regulated, including the fatty acids oleate, linolelaidate, and eicosadienoate. Strikingly, MIE again displayed a delayed response, with 31 differentially regulated metabolites at 0h, increasing to 183 metabolites at 3h. Most of the differentially regulated metabolites were fatty acids, including acetate, beta-hydroxybutyrate, oleate, eicosadienoate, and hexadecadienoate (Figure 2C). Supporting these data, cluster-based analysis of all differentially regulated metabolites revealed 5 distinct expression patterns, most demonstrating time-dependent changes following SIE (Figure 2D).

**Figure 2.**
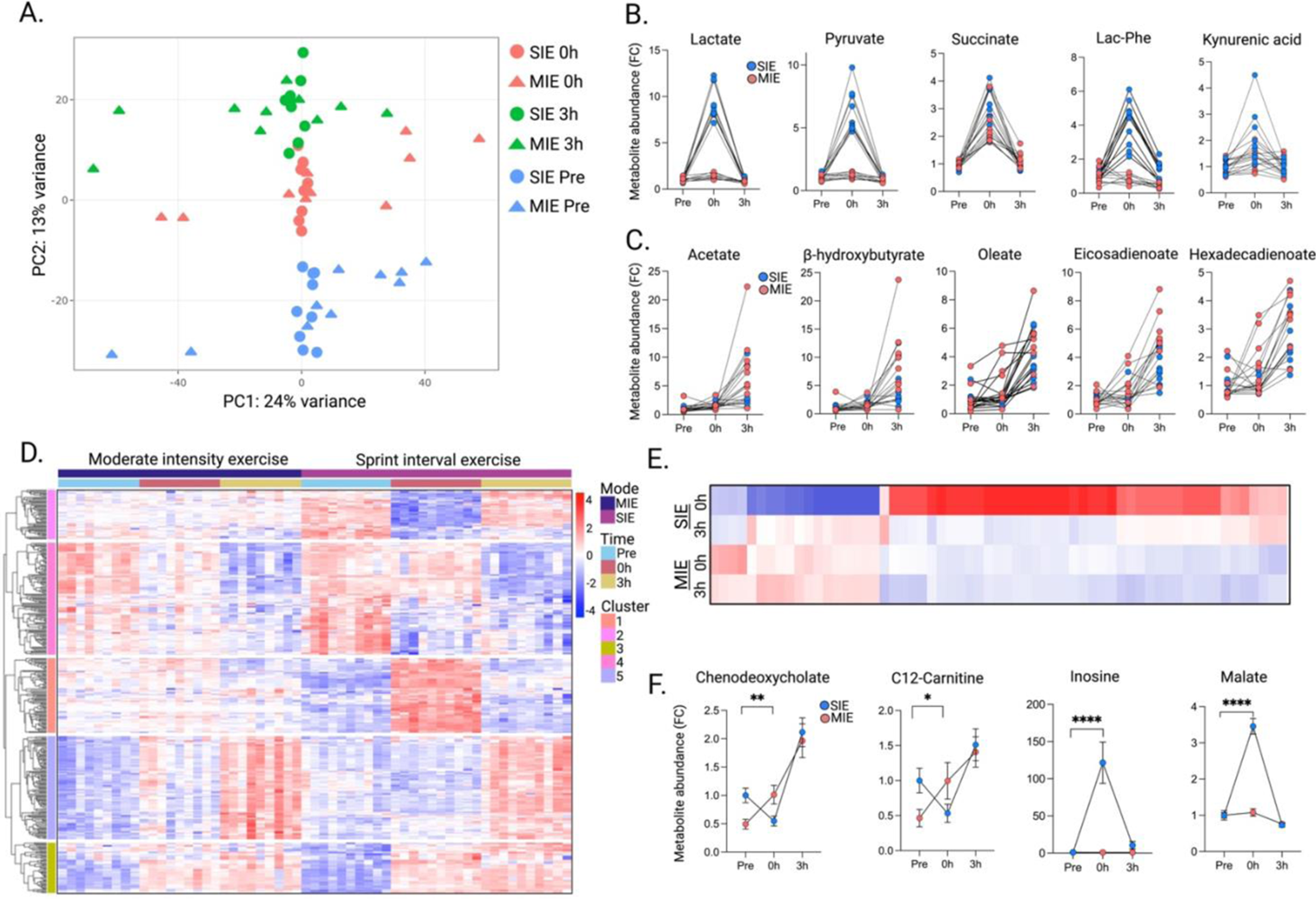
The plasma metabolome following SIE and MIE. **A.** Principal component analysis of individual samples before (pre), immediately following (0h), or 3-hours following (3h) SIE or MIE. Line plots showing representative metabolites which peak at either 0h (**B.**) or 3h (**C.**). **D**. A heatmap showing all differentially regulated metabolites, and how they cluster by temporal expression patterns. **E**. A heatmap of the intensity-dependent differentially regulated metabolites. **F**. Representative metabolites which are differentially regulated between SIE and MIE. Error bars indicate standard error of the mean. \**p*<0.05, \*\**p*<0.01, \*\*\*\**p*<0.0001.

Characterizing metabolites differentially regulated between exercise intensities uncovered the greatest differences at the 0h timepoint, with nearly all metabolites increasing following SIE compared to MIE (Figure 2E). These included lactate, pyruvate, malate, and inosine — emphasizing elevated glycolytic, TCA, and ATP turnover (Figure 2F). Despite relatively few changes immediately following MIE, we observed fatty acid-derived metabolites and bile acids, including C12-carnitine and chenodeoxycholate, increased immediately following MIE relative to SIE (Figure 2F). These MIE-specific changes could reflect intensity-dependent effects on liver metabolism, a key source of bile acids^27^. Collectively, the plasma metabolome displayed similar quantitative and temporal changes to the plasma proteome, further highlighting intensity-dependent changes to blood-borne factors.

### Multiple organs are predicted to contribute to the plasma proteome following exercise

We next sought to identify the predicted tissue source(s) of exercise-responsive analytes, with a particular emphasis on proteins due to the challenging nature of identifying the cellular origin of circulating metabolites. To this end, we leveraged bulk human RNA-sequencing datasets of diverse tissues from the Genotype-Tissue Expression (GTEx) repository^28^. We considered a gene as enriched within a given tissue if its expression in that tissue was at least four-times higher than its average expression in all other tissues^29^. Using these criteria to filter the differentially regulated plasma proteins following SIE, we found immune cell-enriched proteins changed the most at both 0h and 3h, including AZU1, MNDA, and RAB44. Additional tissue-enriched proteins following SIE were CELA2A (pancreas), PRL (pituitary), TFF2 (stomach), GUCA2A (intestine), CFHR5 (liver), PTH (parathyroid), POMC (brain), EGLN1 (muscle), and FABP4 (adipose) (Figure 3A, 3C, and S4A). Similar to SIE, immune cell-enriched proteins changed the most following MIE, again including AZU1 and RAB44. There was a greater distribution of organ-enriched proteins 3h following MIE, including IGFBP1 (liver), PLIN1 (adipose), PRL (pituitary), GCG (pancreas), and DUSP13A (muscle) (Figure 3B, 3C, and S4B). Critically, cross-referencing our tissue-enriched genes with a multi-tissue human proteomic dataset^30^, strongly corresponded with our gene-centric approach, further emphasizing diverse interorgan signaling following exercise.

**Figure 3.**
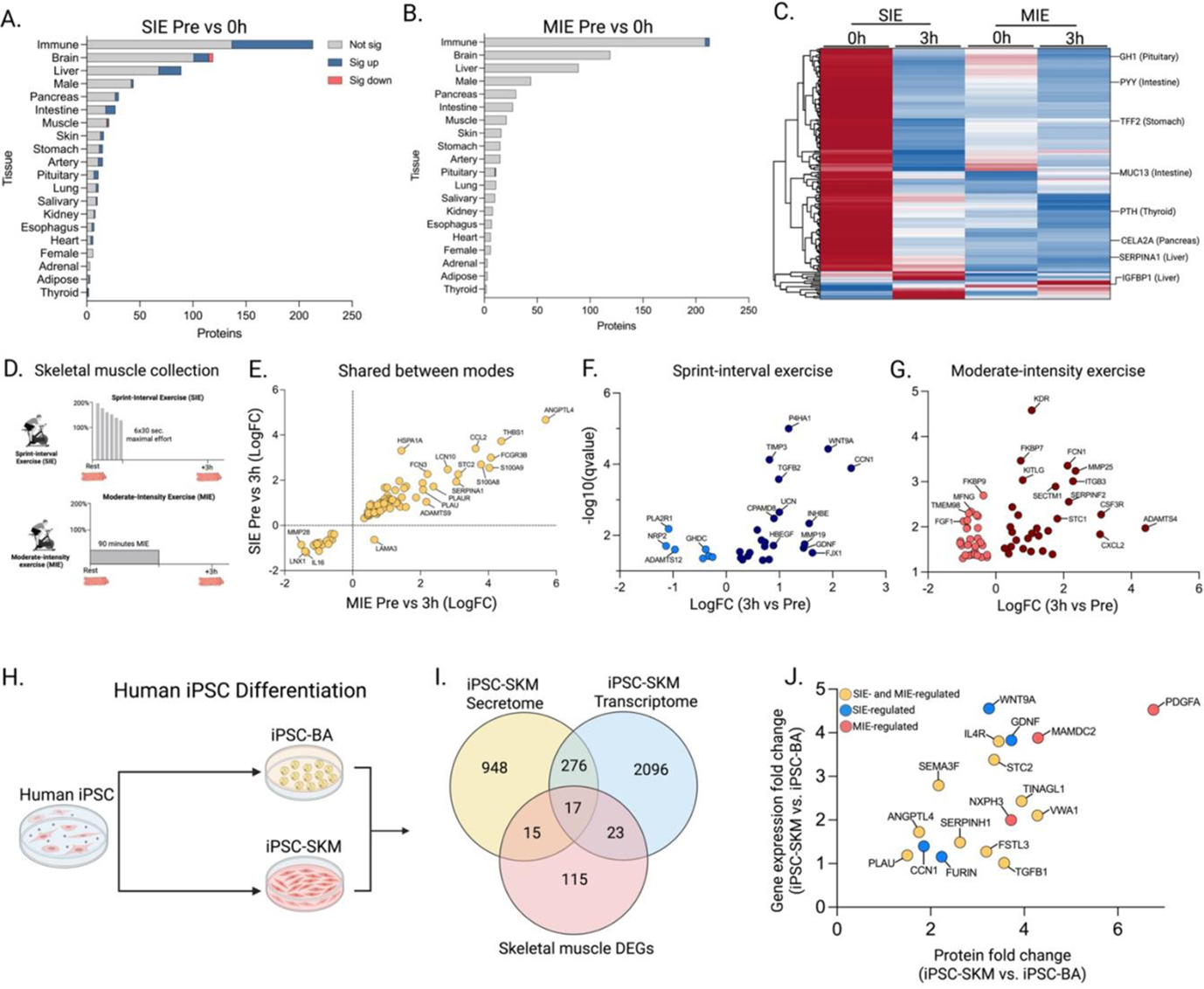
Identifying the tissue of origin for exercise-regulated plasma proteins. **A**. All organ-enriched proteins with those increased (blue bar), decreased (red bar), or no change (grey bar) at 0h following SIE. **B**. All organ-enriched proteins with those increased (blue bar), decreased (red bar), or no change (grey bar) at 0h following MIE **C.** A heatmap showing all organ-enriched proteins differentially regulated at 0h or 3h following SIE and MIE. **D**. A schematic of the muscle collection timepoints following SIE and MIE. Predicted muscle-secretory genes differentially expressed following both MIE and SIE (**E.**), SIE-only (**F**.), or MIE-only (**G.**). **H.** A schematic of human iPSC differentiation into brown adipocytes (iPSC-BA) or skeletal muscle cells (iPSC-SKM). **I**. Venn diagram of the differentially expressed genes and proteins in iPSC-SKM relative to iPSC-BA, and the differentially expressed genes in skeletal muscle following either MIE or SIE which are predicted to encode secreted proteins. **J.** Scatter plot showing the gene expression and protein expression fold change between iPSC-SKM relative to iPSC-BA for the seventeen shared genes/proteins from Figure 3I. The color scheme corresponds with Figure 3E-G; Yellow: shared, Blue: SIE-specific, Red: MIE-specific.

To more directly identify tissue-derived proteins potentially missed using the above-described approach, we collected skeletal muscle from the *vastus lateralis* before and 3 hours following SIE and MIE, and performed bulk RNA-sequencing (Figure 3D)^17^. We chose skeletal muscle because of its documented intensity-dependent remodeling following exercise^31^ and its role in interorgan crosstalk^32^. To identify genes predicted to encode secretory proteins, we overlayed the differentially expressed genes (DEGs) between SIE and MIE with a curated human secretome database sourced from the Human Protein Atlas^33^. This approach revealed 111 and 140 DEGs following SIE and MIE, respectively, predicted to encode secretory proteins (Table S3). Of these, 81 genes were shared between exercise modes, with 30 genes specific to SIE, and 59 genes specific to MIE. Many of the genes shared between exercise modes encode well-described exerkines, including *ANGPTL4*, *VWA1*, *METRNL*, and *HSPA1A* (Figure 3E). SIE- specific genes included *WNT9A*, *CCN1*, and *TIMP3* (Figure 3F), whereas MIE-specific genes included *APLN*, *FCN1*, and *MMP25* (Figure 3G).

While insightful, bulk RNA-sequencing lacks cell-type resolution, preventing the identification of muscle fiber-derived secretory proteins. To overcome this limitation, we isolated and differentiated human inducible pluripotent stem cells (iPSC) into skeletal muscle cells (iPSC-SKM) or brown adipocytes (iPSC-BA), followed by RNA-sequencing (Figure 3H). We included brown adipocytes due to their shared progenitor with skeletal muscle cells, yet distinct differentiation trajectory, serving as a valuable non-muscle control to identify muscle-enriched gene expression^34^. Critically, among the upregulated genes in iPSC-SKM were canonical muscle markers *MYOD*, *MYOG*, and *MYF5,* confirming their muscle-like phenotype (Figure S4C). We next overlayed the bulk skeletal muscle gene expression datasets with the enriched iPSC-SKM genes. Of the original 170 genes predicted to encode secretory proteins (Figure 3E-G), 40 were significantly increased in expression in iPSC-SKM compared to iPSC-BA (FDR<0.01, Log_2_FC>1; Table S4). To confirm that these genes encode secretory proteins, we quantified their protein expression in the conditioned media using Olink-based proteomics^35^. Of the 40 genes increased in iPSC-SKM, 17 of their encoded proteins were also increased in the conditioned media (FDR<0.01, Log_2_FC>1; Table S5). These included SIE-regulated genes (CCN1, FURIN), MIE- regulated genes (PDGFA, MAMDC2), and genes shared across both exercise modes (ANGPTL4, VWA1, PLAU, and TGFB1) (Figure 3I and 3J). As an orthogonal validation, click-chemistry enrichment of azide-labelled proteins secreted from primary human muscle cells identified many of the same proteins — 46 of the original 170 exercise-regulated genes — including CCN1, MAMDC2, ANGPTL4, METRNL, and TIMP3 (see methods; Figure S4D). Collectively, these data highlight how multiple tissues contribute to acute changes in the plasma proteome in an intensity-dependent manner.

### Transcriptomic remodeling of adipocytes following SIE or MIE

Next, we sought to determine the cellular/tissue targets of these secretory proteins. To do so, we integrated a human receptor database from CellTalkDB that includes all known or predicted ligand-receptor pairs^36^. We filtered this list for tissue-enriched receptors based on GTEx gene expression levels as described above and then queried our plasma protein dataset to identify ligands known or predicted to signal through any of the remaining tissue-enriched receptors. This approach revealed multiple ligand-receptor (Ligand:Receptor) pairs differentially regulated immediately following acute SIE, including interactions with the brain (OMG:LINGO1), immune cells (ANXA1:FPR1), adrenals (POMC:MC2R), intestine (GUCA2A:GUCY2C), and kidney (PTH:PTH1R). Predicted crosstalk persisted, albeit to a lesser degree, 3h following SIE, including with the brain (PENK:OPRK1), immune cells (IL1RN:IL1R2), and adrenals (POMC:MC2R) (Table S6). We did not identify any ligand-receptor pairs following MIE, likely reflecting our stringent inclusion criteria.

To more thoroughly characterize how exercised plasma signals to cells in an intensity-dependent manner, we conducted a media swap experiment in primary human adipocytes, a cell type argued by some^37,38,39,40^, though not all^41^, to undergo metabolic remodeling according to exercise intensity. To interrogate this, *in vitro* differentiated human subcutaneous adipocytes were incubated for 12 hours with plasma collected either before or immediately following SIE or MIE from 5-6 subjects per exercise type, followed by bulk RNA-sequencing (Figure 4A). Principal component analysis revealed clear intensity-dependent clustering of the adipocyte transcriptome, with SIE-treated adipocytes shifting along PC1 and little change in MIE-treated adipocytes (Figure 4B). Supporting this, SIE-treated adipocytes displayed robust transcriptomic changes, with 3,560 and 2,586 up- and down-regulated genes, respectively (FDR<0.05, Figure 4C). These included metabolic regulators *VEGFA*, *NR4A3*, *NAMPT*, and *SIRT1* (Figure 4D). In contrast, MIE-treated adipocytes had 369 and 70 genes up- and down-regulated, respectively (FDR<0.05, Figure 4C); most of these were also differentially expressed in SIE-treated adipocytes (346 of 439) such as *CISH* and *APOL4*. Gene set enrichment analysis revealed multiple metabolic pathways enriched within SIE-treated adipocytes, including “rhythmic process”, “organic acid catabolic process”, and “lymphocyte differentiation” (Figure 4E). Supporting these data, transcription factor prediction analysis identified pathways regulated by ARNT and SREBP1 as the most upregulated following SIE plasma treatment (Figure S5A). MIE- treated adipocytes were similarly enriched for metabolic pathways, such as positive enrichment for “cellular response to hormone stimulus” and “chromatin organization”. Surprisingly however, most enriched pathways in MIE-treated adipocytes were negatively regulated, including “electron transport chain”, “oxidative phosphorylation”, and “aerobic respiration” (Figure 4E) — a seemingly counterintuitive response to the aerobic-dependent mode of exercise.

**Figure 4.**
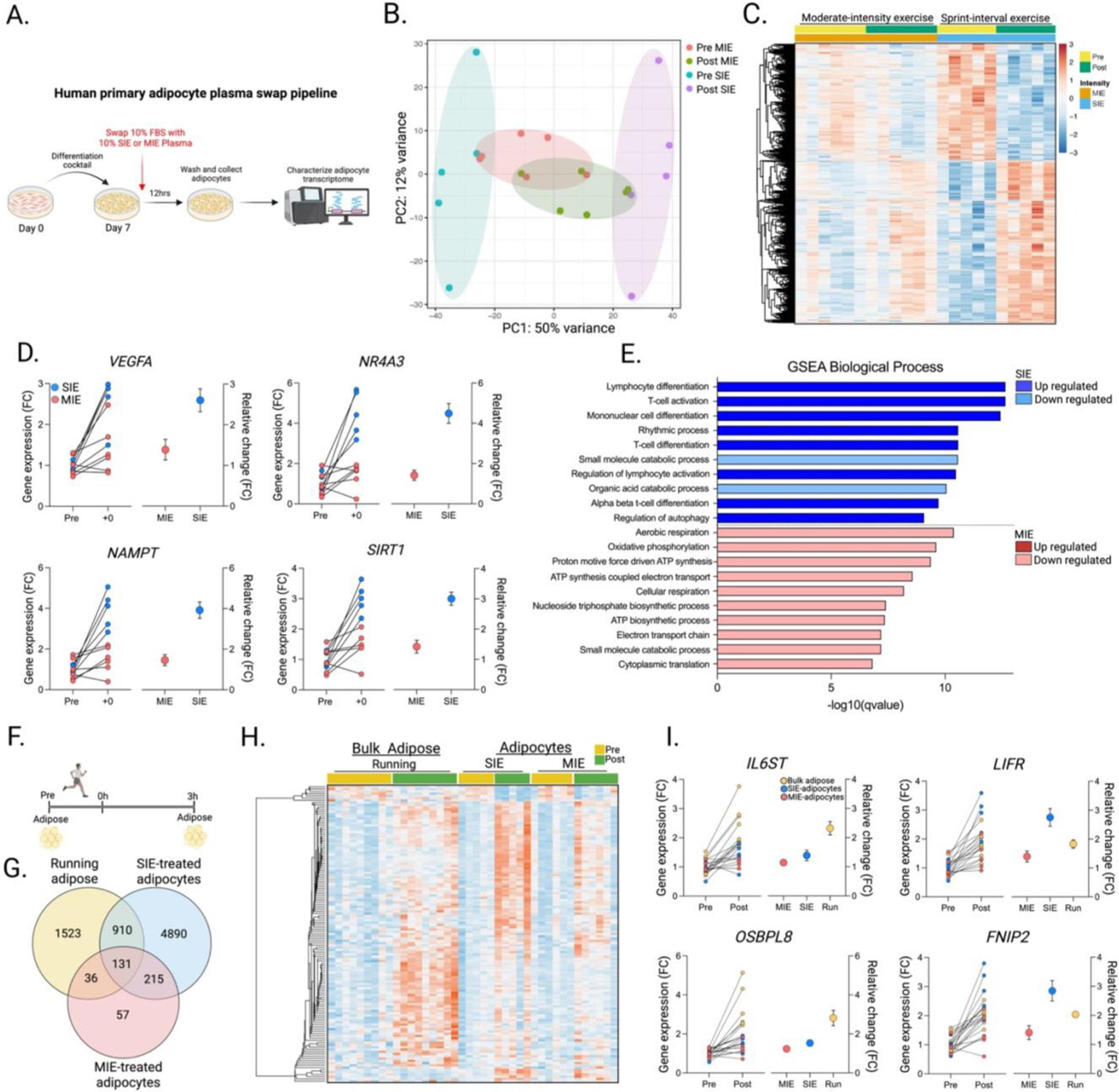
Transcriptomic remodeling of adipocytes following SIE or MIE. **A**. Experimental design of the plasma swap with primary human white adipocytes. **B**. Principal component analysis of the adipocyte transcriptome with pre or 0h plasma from SIE or MIE participants. 5-6 plasma samples from individual participants per mode were used. **C**. Heatmap of all differentially expressed genes following SIE or MIE plasma. **D**. Adipocyte genes of interest and their fold change – including individual values and mean fold change across groups – following incubation with SIE or MIE plasma. **E**. Gene Set Enrichment Analysis (GSEA) of Biological Pathways for all detected genes ranked by change in expression followed by SIE or MIE plasma-treated adipocytes. **F**. Exercise schematic of adipose tissue collected before (pre) and three-hours (3h) following a graded treadmill exercise test. **G**. Venn diagram of the DEGs from the plasma swap samples (Figure 4C) and those following treadmill exercise. **H**. Heatmap of all shared DEGs from Figure 4G. **I**. Genes of interest (fold change and mean fold change across groups) displaying similar expression profiles across datasets.

Despite these intriguing observations, very high-intensity exercise has been shown to decrease adipose tissue blood flow, potentially diminishing acute exposure to the systemic circulation^42^. To confirm that the above-described findings persist within intact adipose tissue, we conducted a maximal graded treadmill exercise test in nine young (mean age = 28.3 years), metabolically healthy male and female participants, and collected subcutaneous abdominal adipose tissue before and 3 hours following exercise. We used treadmill exercise to maximize body-wide metabolic changes, specifically of adipose tissue^43^ (Figure 4F). Bulk RNA-sequencing of adipose tissue revealed clear exercise-dependent clustering (Figure S5B), with 2,600 genes differentially expressed 3 hours following exercise, including 1,429 and 1,171 genes increased and decreased, respectively (FDR<0.05; Figure S5C). The most differentially regulated genes following exercise were *HIPK2* and *ZBED6*, in addition to well-described exercise-regulated genes *ANGPTL4* and *DICER1*. Pathway analysis of upregulated genes again highlighted elevated cellular response to a hormone stimulus, in addition to multiple lipid biosynthetic signaling pathways (Figure S5D)

To identify shared differentially expressed genes within our *in vitro* and *in vivo* datasets — indicating cellular signaling driven, at least in part, by circulating factors — we overlayed the DEGs between SIE-treated adipocytes, MIE-treated adipocytes, and bulk adipose tissue. As anticipated, most shared DEGs were between SIE-treated adipocytes and bulk human adipose tissue. However, 131 genes overlapped between all three datasets (Figure 4G and 4H; Table S7). These included the lipid droplet binding protein *OSBPL8*, the energy sensor *FNIP2*, and receptors regulating cytokine signaling *LIFR* and *IL6ST* (Figure 4I). Intriguingly, multiple adipocyte receptors and their cognate ligands were expressed concordantly across transcriptomic and plasma proteomic datasets, most notably those in the IL6 cytokine family, showing greater expression following SIE. These findings highlight potential intensity-dependent adipocyte regulatory nodes. Collectively, these data underscore plasma-dependent changes to adipocytes *in vitro*, with many shared features in bulk human adipose tissue, and underscore marked cellular rewiring in response to circulatory factors in an intensity-dependent manner.

## Discussion

A single session of exercise stimulates a whole-body metabolic response to meet increasing cellular energetic demands. Interorgan crosstalk plays a critical role in communicating these dynamic metabolic states through the release of metabolites, peptides, proteins, and extracellular vesicles^44^. Soluble proteins have classically been characterized for their role in regulating interorgan crosstalk, yet the broad dynamic range of protein concentrations in the blood — spanning 10 orders of magnitude — has hampered discovery efforts. Moreover, how different exercise modes (cycling versus running), intensities (high-intensity versus low-intensity), and training status (untrained versus trained) uniquely influence interorgan crosstalk in humans has remained understudied.

Employing a multiplexed, antibody-based protein detection platform, we detected nearly 3,000 proteins across more than 100 plasma samples, identifying marked intensity-dependent changes to the plasma proteome. Our findings highlight the dramatic metabolic stimulus provided by sprint-interval exercise, which altered ∼25% of the detected plasma proteins. These changes were observed in both trained and untrained participants, suggesting protein release through cellular leakage is not a primary source. Nonetheless, the presence of cytosolic proteins that are not considered secretory — such as the kinase NADK — may reflect increased cellular turnover following exercise. Despite this, multiple signaling proteins were increased and decreased following SIE to a greater degree than MIE, including those regulating angiogenesis (VWF), extracellular matrix remodeling (TIMP3), gut signaling (TFF2), and potential neuro-regulation (POMC). In addition, plasma metabolomics similarly reflected the acute energetic demands stimulated by SIE, with plasma lactate, pyruvate, malate, and fumarate increased immediately following exercise. In contrast, fatty acids displayed a delayed release into the plasma, likely underscoring the shift from carbohydrate metabolism toward lipid metabolism during recovery.

Although MIE produced only modest overall changes to the plasma proteome, the proteins that did change are well-described endocrine factors such as ANGPTL4, IGFBP1, and FST — the latter two increasing exclusively following MIE. These MIE-dependent responses may reflect the sustained energy demands required by continuous exercise, most notably an increase in the glucagon-to-insulin ratio and liver glycogen depletion, which stimulates hepatic secretion of FST and IGFBP1^45,46,47^. Indeed, the liver — in addition to adipose tissue — may be uniquely sensitive to exercise intensity and duration, as many of the most differentially regulated plasma proteins at 3h are hepatokines^46^. Additionally, the temporal changes to the plasma proteome could reflect reliance on transcription-driven changes to secretory proteins that are stimulated during the MIE bout, with their protein abundance and subsequent secretion, peaking at 3h. Intriguingly, the delayed secretory kinetics were not unique to proteins, but also observed with metabolites, most notably increased fatty acids at 3h. These temporal patterns following MIE likely reflect both intensity-dependent and volume-dependent changes to the secretome — though we were unable to address the latter variable in our exercise paradigm.

A potentially underappreciated mode of cellular crosstalk during exercise, specifically very high-intensity exercise, is ectodomain shedding of transmembrane proteins. These proteolytically cleaved transmembrane proteins contribute to the “sheddome”^48^, and regulate multiple physiological processes including skeletal muscle, neural, and liver metabolism^49,50,51^. In fact, recent findings revealed the hepatokine Syndecan-4 (SDC4) undergoes ectodomain shedding in response to exercise, stimulating improved hepatic metabolic function^51^. While our Olink-based approach is unable to distinguish ectopically cleaved proteins, the increased abundance of multiple metalloproteases — proteases contributing to ectopic protein cleavage — supports the sheddome as a relevant mode of interorgan crosstalk following exercise.

The identification of multiple intensity-dependent proteins within the circulation suggests dynamic interorgan crosstalk. Using a previously-validated protocol leveraging the GTEx gene expression resource^28,29^, we found several organs contribute to the plasma proteome, with more diverse organ contribution following SIE. To more directly address organ contribution, we investigated skeletal muscle, an organ that contributes to systemic metabolism through myokine secretion and interorgan crosstalk^32^. We characterized bulk skeletal muscle, iPSC-derived human muscle cells, and primary human muscle cells to characterize the expression and secretion of proteins from myofibers following exercise. While we detected multiple differentially regulated genes following SIE or MIE in bulk skeletal muscle tissue, and confirmed their ability to be secreted from either iPSC-derived or primary human muscle cells, there was minimal concordance between the differentially regulated plasma proteins and our muscle datasets. In fact, several predicted secretory proteins that increased in both gene expression and protein secretion *in vitro* were unchanged in the plasma. Those that did change, such as ANGPTL4, do not have a net release from the skeletal muscle in response to exercise^52^. These findings are supported by recent cell type-selective tagging of secreted proteins following exercise in mice, detecting only minimal muscle fiber-derived proteins in the blood^53^. Intriguingly, however, robust secretion is observed in the skeletal muscle interstitium^54,55^. These data suggest preferential autocrine and paracrine signaling in human skeletal muscle following exercise, with relatively few myokines (e.g., IL-6^56^) entering the systemic circulation. More work will be required to address this.

Our data highlights the sensitivity of primary human adipocytes to plasma collected following SIE or MIE. We identified striking remodeling of the adipocyte transcriptome following treatment with SIE plasma, stimulating a nearly 10-fold increase in the number of differentially expressed genes compared to MIE. These findings were particularly surprising since low- and moderate- intensity exercise preferentially rely on lipid metabolism, with high- and very high-intensity exercise relying on carbohydrate metabolism^57,58^. In fact, blood flow to the subcutaneous adipose tissue has been shown to decrease during very high-intensity exercise^42^. Nonetheless, glycerol release into the circulation increases following very-high intensity exercise, indicating adipocyte metabolism is modified by sprint-interval exercise. To address this, we integrated the *in vitro* adipocyte gene expression datasets with bulk human subcutaneous adipose tissue collected following a treadmill exercise test to exhaustion. While we did observe gene expression overlap, it was relatively minor (∼5% of the transcriptome). This could reflect multiple cell populations in bulk adipose tissue, potentially diluting the adipocyte gene expression signal. Genes that did overlap across datasets included membrane-bound receptors, including those from the IL6 cytokine family *IL6ST* and *LIFR*. Intriguingly, ligands for these receptors were similarly increased in the plasma proteome, most notably oncostatin-M (OSM) and IL6. We previously found OSM is a macrophage-derived ligand signaling to adipocytes following exercise^59^. Moreover, while our data suggest intensity-dependent regulation of adipocytes derived from the abdominal subcutaneous depot, it does not reflect depot-specificity, such as intermuscular adipocytes, which are dynamically responsive to metabolic perturbations^60,61,62^. Future work will be necessary to determine whether these intensity-dependent changes across adipose tissue depots persist in humans undergoing different exercise modes and intensities.

Collectively, these data support intensity-dependent changes to the secretome, uncovering distinct shifts in protein and metabolite secretory dynamics, and identify their predicted tissue of origin and target tissue. Despite the need for future studies with larger sample sizes, more sample collection timepoints, and mechanistic characterization of our detected secretory proteins, these data nonetheless offer new insights into how distinct exercise intensities and modes influence systemic metabolism in humans. While further work is required to confirm their metabolic significance, our findings provide a new foundation from which future research can dissect the functional significance of exercise-regulated secretory proteins and metabolites in human health.

## Methods

### Human exercise studies: Cohorts 1-3

#### Cohort 1

Plasma samples were obtained from Botella et al. 2024^17^. Ethical approval was obtained from the Victoria University Human Research Ethics Committee (HRE17-075) and registered as clinical trial ANZCTR; ACTRN12617001105336. Of the original 28 subjects recruited, 19 were used for this study due to sample availability. Detailed methods for the exercise intervention and sample collection can be found in Botella et al. 2024^17^. In brief, 28 healthy males volunteered to participate in acute exercise testing before and after an 8-week training intervention. Participants were matched based on their pre-determined maximum aerobic power followed by random assignment to either sprint-interval exercise or moderate intensity exercise. Participants were provided a balanced set of meals 24 hours prior to performing a single exercise session. The sprint interval exercise session consisted of six, 30-second all-out cycling bouts against a resistance set at 0.075 kg•kg BM-1, interspersed by 4-minute recovery periods. Sprint interval training comprised 3-4 sprint interval exercise sessions per week, with the resistance gradually increasing from 0.075 kg•kg BM-1 to 0.090 kg•kg BM-1 at week seven. Exercise sessions increased to eight sprints per session by week seven. The moderate-intensity exercise session consisted of continuous cycling at a fixed power equivalent to ∼ 90 to 100% of the predetermined first lactate threshold for 90 minutes. Moderate-intensity training consisted of progressively increasing exercise duration to 120 minutes per session in week seven, with the intensity adjusted based on submaximal tests performed every two weeks. Blood samples were collected before, immediately following, and 3 hours following the single exercise sessions in the untrained and trained state.

#### Cohort 2

Plasma samples were obtained from Axelrod et al. 2019^19^ (IRB#: 15-1311) with detailed methods described in the original publication. In brief, eleven active, young, male participants arrived for testing in a fasted state at 0600, and received a standardized meal one hour prior to exercise testing. Exercise consisted of two hours of continuous running on a motorized treadmill at 60% of their pre-determined VO_2max_. Blood was collected before and immediately following exercise.

#### Cohort 3

This study was approved by the Rockefeller University Institutional Review Board and registered as a clinical trial (IRB #: LOL1044; Clinical trial ID: NCT06223035). A total of eleven healthy, moderately active participants (N=7 biological males; N=4 biological females) were recruited to the Rockefeller University Hospital, provided their written consent, and completed all study sessions. The protocol consisted of two in-person visits. Visit one consisted of participants arriving at the hospital in the morning (0800-1000). All participants were asked to eat their regular breakfast at least two hours before arriving to the hospital. Body composition measurements were taken using air displacement plethysmography (BodPod; COSMED), and blood draws were taken from the median cubital vein and collected in K2 EDTA-coated tubes for clinical blood work. Following blood collection, participants were prepped for subcutaneous adipose collection from the lower abdomen. The area for tissue collection was numbed with a local anesthetic (lidocaine) and ∼50 mg of adipose tissue was collected using a 6 mm punch biopsy needle. Skin and connective tissue was carefully removed and tissue was blotted for blood. Adipose samples were then immediately frozen in liquid nitrogen and stored at -80 °C for future use.

Following a minimum of one week from adipose sample collection, participants arrived in the morning (0800-1000) to the Sports Rehabilitation and Performance Center at the Hospital for Special Surgery (HSS) for VO_2max_ testing. Participants were asked to consume the same breakfast as they did prior to visit one, at least two hours before arriving to HSS. Following familiarization with the protocol and equipment, participants underwent a five-minute warm up, comprising walking on a treadmill at a slow pace. Following five minutes, participants began the graded exercise test which included a progressive increase in running speed and treadmill incline until participants reached volitional fatigue as defined by either reaching a rate of perceived exertion (RPE) greater than 19 (using the Borg RPE Scale), a respiratory exchange ratio (RER) reaching above 1.1, or if the participant chose to stop the test for any reason. Blood lactate levels were measured before and immediately following exercise via a finger stick. Immediately following the exercise test, participants returned to the Rockefeller University Hospital for subsequent blood and adipose collection. Three hours following the exercise test, participants were prepped for abdominal subcutaneous adipose collection as described above. The opposite side of the abdomen (left or right of the navel from which the first biopsy was collected) was used for the second adipose collection. Upon collection, adipose was cleared of remaining skin and connective tissue, blotted for blood, and rapidly frozen. One male participant chose not to complete the second adipose biopsy and was thus not included. Additionally, the adipose sample from one female participant became compromised due to accidental exposure to room temperature and was not included. The final sample size for RNA-sequencing was N=9 (N=6 biological males; N=3 biological females). Frozen adipose tissue was homogenized in a bead beater with Trizol (ThermoFisher Scientific: 15596026). RNA was separated via the Trizol-chloroform method. The subsequent RNA fraction was combined with 70% ethanol, which was then processed with the Qiagen RNeasy Plus Micro Kit (Qiagen: 74034), including DNAse incubation treatment. High-quality RNA was processed for RNA-sequencing as described in the RNA-sequencing section.

### Sample processing for plasma OLINK proteomics

Plasma samples from Cohort 1 and Cohort 2 underwent proteomic profiling in a single batch using Olink® Explore 3072 panel (Olink Proteomics AB, Uppsala, Sweden) according to the manufacturer’s instructions; a detailed description of methods can be found elsewhere^63^. In brief, the Olink protocol is based on Proximity Extension Assay (PEA)^64^, coupled with readout via next-generation sequencing (NGS) and enables the detection of ∼3,000 proteins within a given biological sample. Specifically, pairs of oligonucleotide-labeled antibodies bind to their target protein, followed by nucleotide hybridization, polymerization, extension, and amplification using NGS Illumina® NovaSeq™ 6000. Critically, these steps only occur if the oligonucleotides are in close proximity, decreasing the likelihood of non-specific protein detection. The raw data output is quality controlled, normalized, and converted into NPX® values, Olink’s proprietary unit of relative abundance. All assay validation data (detection limits, intra- and inter-assay precision data) are available on manufacturer’s website (www.olink.com/explore).

### Sample processing for plasma untargeted metabolomics

Samples (20 µL) were extracted with 80 µL of 75:25:0.2 ACN:MeOH:FA extraction solution containing 1 µM heavy amino acid mix. After vortexing and centrifuging (at 13,200 rpm and 4 °C for 15 min), 40 µL supernatant was transferred to HPLC vials. The remaining supernatants were combined to prepare a pooled QC sample to monitor for instrument variability such as mass error, chromatographic shift, and internal standard signal. Samples were analyzed using LC-MS on an IQ-X Orbitrap mass spectrometer coupled with a Vanquish UPLC System (ThermoFisher Scientific). For chromatographic separation, samples were loaded on a SeQuant ZIC-pHILIC column (150 × 2.1 mm, 5 μm polymeric) and a SeQuant ZIC-pHILIC Guard Kit (20 × 2.1 mm) at 40 °C and eluted with a solvent system composed of mobile phase A (20 mM ammonium carbonate and 0.1% ammonium hydroxide in water at pH 9.3) and mobile phase B (100% acetonitrile). The injection volume was set to 5 µl, and samples were maintained at 4 °C. The gradient (vol/vol) used was as follows: 0–22 min, linear gradient from 90% to 40% B; 22–24 min, held at 40% B; 24–24.1 min, returned to 90% B; 24.1–30 min, equilibrated at 90% B at a flow rate of 150 µl/min. Mass spectrometry was operated in polarity-switching mode for MS1 acquisition with the following source conditions: spray voltage, +3 kV and -2.5 kV; sheath gas, 40 AU; auxiliary gas, 15 AU; ion-transfer tube temperature, and probe heater temperature, 300°C. Samples were analyzed with the following acquisition parameters. MS1 scans: resolution, 120,000; AGC target, 1.2 × 10^5^; maximum injection time, 246 ms; m/z ranges, 55-275 Th and 265-1000 Th. The pooled QC sample was analyzed using data-dependent acquisition method. Data-dependent MS/MS scans were acquired separately for each polarity (positive and negative) and mass range (55-275 Th and 265-1000 Th) using the following settings: a resolution of 15,000, 7.5e4 AGC target, auto max injection time mode, 1 Da isolation width, HCD activation, assisted normalized collision energy of 20, 30, 40 units, and a cycle time of 3 seconds. The instrument was externally calibrated weekly using Pierce FlexMix Calibration Solution (ThermoFisher Scientific) and the internal calibration feature (RunStart EASY-IC) was turned on.

### Differential analysis of the Olink plasma proteome and untargeted metabolites

For the proteome, the mean Olink NPX value for each protein was calculated and missing values were imputed using the DEP R Bioconductor package (v1.26.0). Specifically, imputation was done with the impute function with the ‘knn’ function and the ‘rowmax’ parameter set to 0.9. For both the proteome and metabolome, differential abundance between time points within each mode of exercise (sprint interval vs moderate) was determined using the limma R Bioconductor package (v3.60.4). The correlation of abundance from the same subject across time points was estimated using the duplicateCorrelation function from limma, and this was included in the linear model fitting function (lmFit) with the ‘block’ parameter when comparing time points within each mode of intensity. Moderated t-statistics were obtained using empirical Bayes, and the topTable functions implemented within limma was used to identify all proteins that changed across the various timepoints for either mode of intensity after adjusting p-values with the Benjamini– Hochberg procedure. Adjustment for subject effects for visualization was performed using the removeBatchEffect function in limma. The z-scores of these subject-corrected abundance values for differential proteins or differential metabolites was visualized using the pheatmap R package (1.0.12)^65^.

### Azide-tagging of nascent secreted proteins in primary human muscle cells

Human primary myoblasts were obtained from muscle biopsies taken from the *vastus lateralis* of healthy volunteers^66^. Cells were left to proliferate in growth media consisting of DMEM/F12+Glutamax (Gibco: 31331), 1% Antibiotic-Antimycotic (Anti-anti) (Gibco: 15240-062), and 16% FBS (Sigma: F7524). Upon confluence, cells were differentiated in media consisting of DMEM (Gibco: 31966-021), 20% Medium 199 (Gibco: 31150-022), 2% HEPES (Gibco: 15630-056), 1% Anti-anti, 0.03 µg/mL Zinc sulfate (Sigma: Z4750), 1.4 µg/ml Vitamin B12 (Sigma: V6629), 2% FBS, 10 µg/mL insulin, and 100 µg/mL apo-transferrin (BBI Solutions: T100-5). Once fully differentiated (minimum of 75% differentiation), cells were washed with warmed PBS and provided fresh DMEM and differentiation cocktail with dialyzed FBS containing either azidohomoalanine (AHA, 0.1 mM; Click Chemistry Tools: 1066) or Methionine (MET, 0.1 mM; Sigma: M5308) for 15 hours. Following 15 hours, the media was removed and fresh media with AHA (0.1 mM) or MET (0.1 mM) were added. Muscle cells were then left unperturbed for 3-hours, followed by collection of the conditioned media for analysis. Collected conditioned media was passed through a 0.22 µm strainer and concentrated (dialyzed) using the amicon 3 kDa filters (Millipore: UFC900308). The concentrated media was placed at -80 °C for long-term storage. Protein concentration was determined using the Pierce BCA protein assay Kit (ThermoFisher Scientific: 23225).

### Click-chemistry enrichment of muscle cell-derived azide-labeled proteins

The overarching principal of Bio-Orthogonal Non-Canonical Amino Acid Tagging (BONCAT) relies on the incorporation of amino acid analogs containing a functional group, specifically an azide, into the cellular proteome. Azidohomoalanine is a methionine analog which differs only in its unique azide moiety. The endogenous protein translation machinery recognizes AHA, and incorporates it into newly synthesized proteins in place of methionine^67^. Azide-bearing proteins— either cytosolic or those secreted into the conditioned media — can undergo affinity purification through introducing an alkyne-bound bead. The alkyne moiety forms a covalent bond with azide when in the presence of a copper catalyst, ultimately ‘clicking’ the azide and alkyne moieties together. As such, this approach only detects proteins synthesized within cells, circumventing the need for serum depletion. We and others have demonstrated the successful use of AHA in primary murine cell lines^68,69^. Our preliminary studies confirmed sufficient incorporation of AHA into the primary myofiber proteome, with 16 hours showing greatest incorporation. All primary muscle cells from each participant successfully incorporated AHA into their respective proteome. Methionine-pulsed samples were used to determine specificity of protein enrichment. 3.5 mg of protein from each sample were used for click enrichment using the Click-iT Protein Enrichment Kit following the manufacturers protocol (Invitrogen: C10416). Following enrichment, protein-bound beads were placed at -80 °C until follow-up proteomic analysis.

### Proteomic processing of bead-bound proteins

Protein-bound beads underwent reduction and alkylation and then on-bead digestion with Trypsin for 3 hours. Supernatant was extracted and proteins were then digested with Lys-C and Trypsin overnight. Digestion reactions were stopped with neat TFA. Samples then underwent Solid Phase Extraction, prior to being analyzed by LCMS (70 min analytical gradient and separated using a 12 cm Pulled Emitter column (C18 reversed phase)). The mass spectrometer (LUMOS) was operated in high res./high mass accuracy mode. Generated data was searched and quantified using ProteomeDiscoverer/Mascot. Data were queried against the human database concatenated with Trypsin and Lys-C sequences. Raw data was processed through MaxQuant (v 2.6.6.0) software using the human database to produce label-free quantification (LFQ) and intensity-based absolute quantification (iBAQ) values. LFQ and iBAQ values were then taken to Perseus to perform statistical analysis. Values were Log2 transformed and filtered to remove proteins that were either found by reverse search or as contaminants. iBAQ values were filtered to only include proteins that produced a value in 4 out of 6 replicates for at least one sample group for AHA vs MET comparisons. “NaN” values were then imputed with random low-abundant signals for statistical analysis purposes. Sample groups (MET vs AHA) were plotted against each other in two-sample t-tests.

### Human skeletal muscle collection and RNA-sequencing

Participant information is described above (Cohort 1). In brief, skeletal muscle was collected under local anesthesia from the *vastus lateralis* using the Bergström needle biopsy technique before and 3 hours following exercise. Muscle tissue was blotted for residual blood and any remaining fat and connective tissue was carefully removed. The sample was then immediately frozen in liquid nitrogen and stored at -80 °C. All RNA extraction, cDNA library, and RNA-sequencing can be found in Botella et al. 2024^17^.

### Human iPSC-derived brown adipocytes and skeletal muscle cells

Human pluripotent stem cell-derived brown adipocytes and skeletal muscle cells were differentiated from NCRM1 (NIH CRM) human iPSC’s as described^70,71^ and is the same sample material used from Plucińska et al. 2025^35^, where detailed cell differentiation and processing is described. Following differentiation of the respective cell-types, media samples were collected from mature skeletal muscle and brown adipocytes and processed for OLINK analysis as described in Plucińska et al. 2025^35^, and were analyzed using the Limma package in R. mRNA from muscle and brown adipocyte cultures were obtained as previously described^35^. mRNA was processed for RNA-sequencing as described below.

### RNA-sequencing

For all RNA-sequencing experiments, 100 ng of total RNA was used to generate RNA-Seq libraries using Illumina stranded mRNA kit (Catalog #: 20040534), following the manufactures protocol. Libraries prepared with unique dual indexes were pooled at equal molar ratios. The pool was sequenced on Illumina NextSeq 2000 P3 flowcell using NextSeq Control Software v1.7.1.46395 to generate 100 bp (or 50 bp) paired reads, following the manufactures protocol (Document #: 200027171 v04)

### Identification of tissue-enriched plasma proteins

To identify proteins in the plasma that were enriched in a particular tissue, bulk RNA raw read counts were downloaded from the GTEx portal (version 8) and normalized using the DESeq2 R Bioconductor package (v 1.44.0)^28,72^. The mean normalized count for each gene within a tissue was calculated, and a gene was determined to be enriched in a particular organ if the maximum expression value for the group of tissues belonging to that organ was four times the value of the maximum of the tissues for next highest organ.

### Prediction of organ-to-organ crosstalk

To predict organ-to-organ crosstalk, we leveraged a human receptor database sourced from CellTalkDB which includes all known or predicted ligand-receptor pairs upon a given cell-type^36^. To filter down the number of receptors, and determine those that are enriched within a given organ, we removed all known or predicted receptors which are not tissue-enriched as determined by their GTEx gene expression levels described above. From the remaining list of receptors, we overlayed the differentially regulated plasma proteins to identify ligands which have a known or predicted receptor from the remaining tissue-enriched receptor dataset.

### Primary human adipocyte media swap with exercised plasma

Abdominal subcutaneous adipose tissue was collected from a healthy, middle aged (47 years) female, and the stromal vascular fraction (SVF) containing preadipocytes were isolated and purchased from Lonza (Catalog #: PT-5001; Lot #: 24TL271510). Preadipocytes were subsequently plated and differentiated into mature adipocytes in 6-well collagen coated plates following the manufacturer’s instructions. Fully differentiated adipocytes were histologically multilocular and positive for adipocyte markers *FABP4*, *FASN*, and *PPARG*, compared to non-differentiated preadipocytes. Upon complete differentiation, adipocytes were washed twice with warm PBS, followed by a 30-minute incubation with 900 µl of Lonza growth media (Catalog #: PT-8002) containing L-glutamine, recombinant human insulin, and gentamicin sulfate-amphotericin, without fetal bovine serum. Following 30 minutes, 100 µl of human plasma was added to each well. Plasma samples were from individuals undergoing either moderate intensity exercise or sprint interval exercise as described in Cohort 1. The resting (pre) or immediately post (0h) plasma samples were used. Each timepoint from each individual were conducted in duplicate. Six participants for either SIE or MIE were used. One plasma sample from SIE resulted in adipocytes becoming detached, and were thus not used for downstream processing. The final sample size was N=6 for MIE and N=5 for SIE. Adipocyte cultures containing plasma were incubated at 37 °C in a humidified incubator with 5% CO_2_ for 12 hours. Following 12 hours, adipocytes were washed with ice-cold PBS and were immediately processed for RNA extraction. In brief, adipocytes were homogenized in Trizol (ThermoFisher Scientific: 15596026), and the RNA was separated via the Trizol-chloroform method. The subsequent RNA fraction was combined with 70% ethanol, which was then processed with the Qiagen RNeasy Plus Micro Kit (Qiagen: 74034) This included DNAse treatment. High-quality RNA was processed for RNA-sequencing as described above.

## Acknowledgements

We would like to thank the Rockefeller University Hospital (RUH), including Lucy Apicello, Rita Devine, Lin Zhen, Regina Butler, Tia Gareau, Jill McCabe, and Andrea Ronning for their help with recruitment and on-site visits. We thank Richard Hutt, Rhonda Kost, Anna Rauso, Vanessa Smith, Dale Miller, and Teresa Solomon for assistance with Institutional Review Board (IRB) submissions, clinical trial submissions, and contract agreements between institutions. We thank Connie Zhao and Bin Zhang from the Genomics Resource Center, and Wei Wang and Thomas Carroll of the Bioinformatics Resource Center for RNA-sequencing, processing, and thoughtful discussions. We thank Elizabeth Zunica for organizing human sample shipments. We thank all participants who volunteered their time to take part in these studies. Figures were created in BioRender (https://BioRender.com).

## Author Contributions

LO and PC conceptualized the study; LO and PC acquired funding; LO and DB analyzed and visualized data; LO, JB, DJB, and PC designed and interpreted experiments; LO, JB, LY, LF, KB, CA, MK, HS, CP, ER, EK, KF, and KP performed experiments; JW collected human adipose tissue samples; PC, DJB, LO, JB, AK, JRZ, EVV, JK and OP supervised study/study segments; REG, JMR, and OP provided technical and intellectual input; LO and PC wrote the manuscript. All authors read and approved the final version of the manuscript.

## Data Availability

All original RNA-sequencing data in this manuscript has been deposited in the Gene Expression Omnibus database (ID: GSE308252). The mass spectrometry proteomics data have been deposited to the ProteomeXchange Consortium via the PRIDE partner repository with the dataset identifier PXD069170. The untargeted metabolomics data will be deposited upon publication.

---

**Figure S1.**
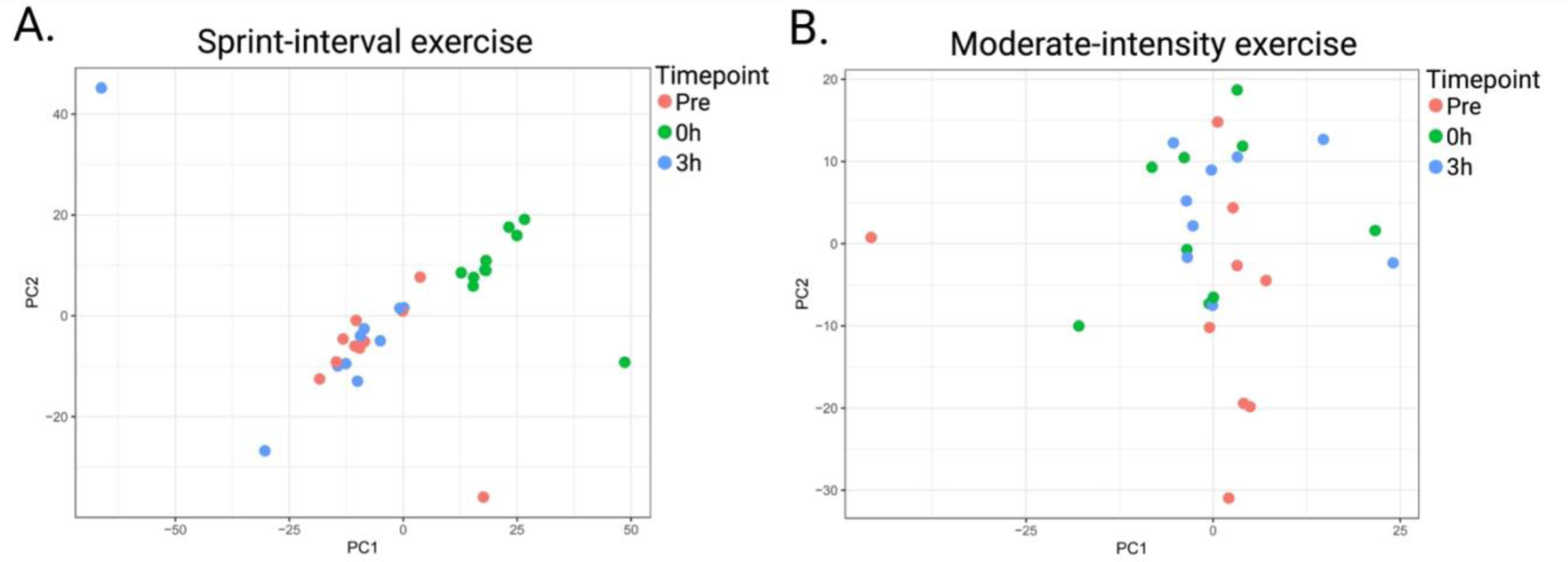
Sample clustering of the plasma proteome following SIE or MIE in the untrained state. Principal component analysis of plasma proteins following SIE (**A**.) or MIE (**B**.) in untrained participants.

**Figure S2.**
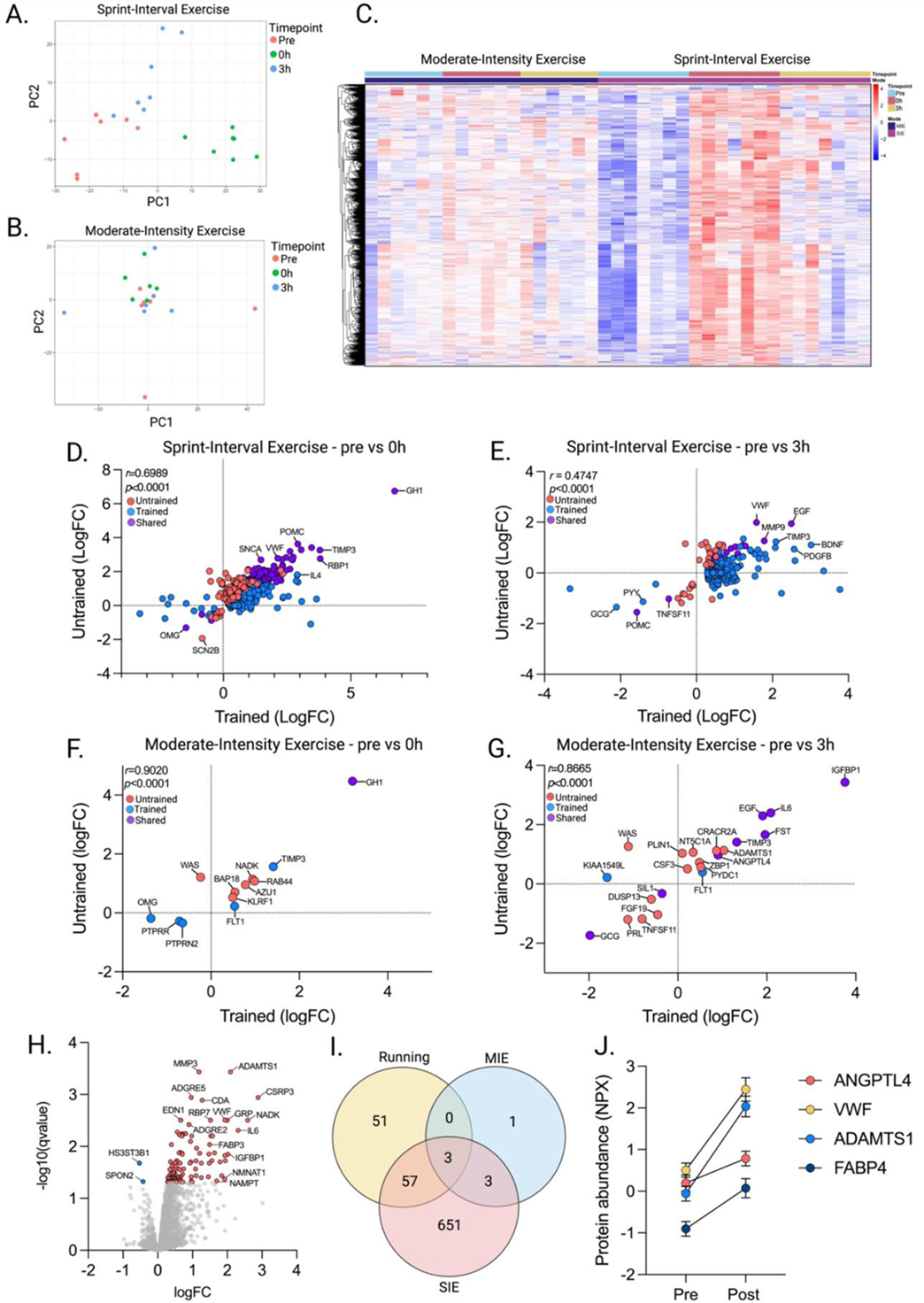
The plasma proteome following SIE or MIE in the trained state, and following moderate intensity running. Principal component analysis of plasma proteins following SIE (**A**.) or MIE (**B**.) in the trained state. **C**. A heatmap of all differentially regulated plasma proteins following a single session of SIE or MIE plasma after 8 weeks of mode-specific training. A plot showing the concordance between untrained and trained samples following SIE at 0h (**D**.), SIE at 3h (**E**.), MIE at 0h (**F**.), and MIE at 3h (**G**.). **H**. Volcano plot showing the differentially regulated plasma proteins before and after moderate intensity running. **I**. Venn diagram of the differentially regulated plasma proteomes immediately following MIE, SIE, and moderate intensity running. **J**. Representative plasma proteins differentially regulated following moderate intensity running. Protein values are Normalized Protein values (NPX). For Figure S1D-S1G, a Pearson’s product-moment correlation (r) to determine linear associations was used. Statistical significance was evaluated with two-tailed tests at *p*=0.05. Error bars indicate standard error of the mean.

**Figure S3.**
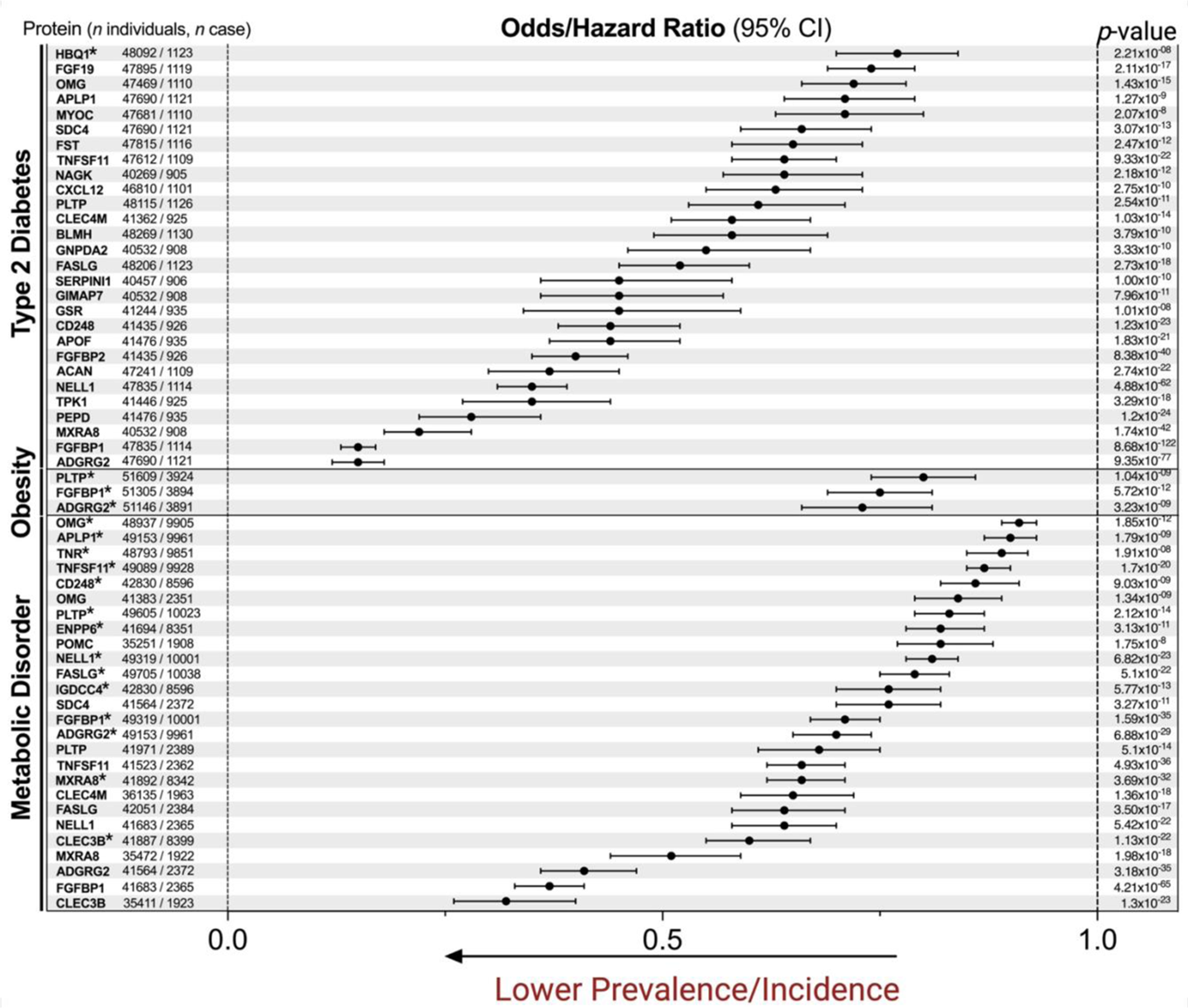
Exercise-regulated proteins are associated with health and disease. Data shown are proteins differentially regulated following SIE or MIE which are significantly associated with either “Metabolic Disorder”, “Obesity”, and/or “Type 2 Diabetes”. Data include “prevalence” and “incidence” and thus either Odds Ratio or Hazard Ratio are plotted. Because a protein can be associated with both the incidence and prevalence for a given trait, some proteins are plotted twice within their respective trait. The asterisk indicates proteins associated with the incidence of a given disease whereas all unmarked proteins correspond to the prevalence of a given disease.

**Figure S4.**
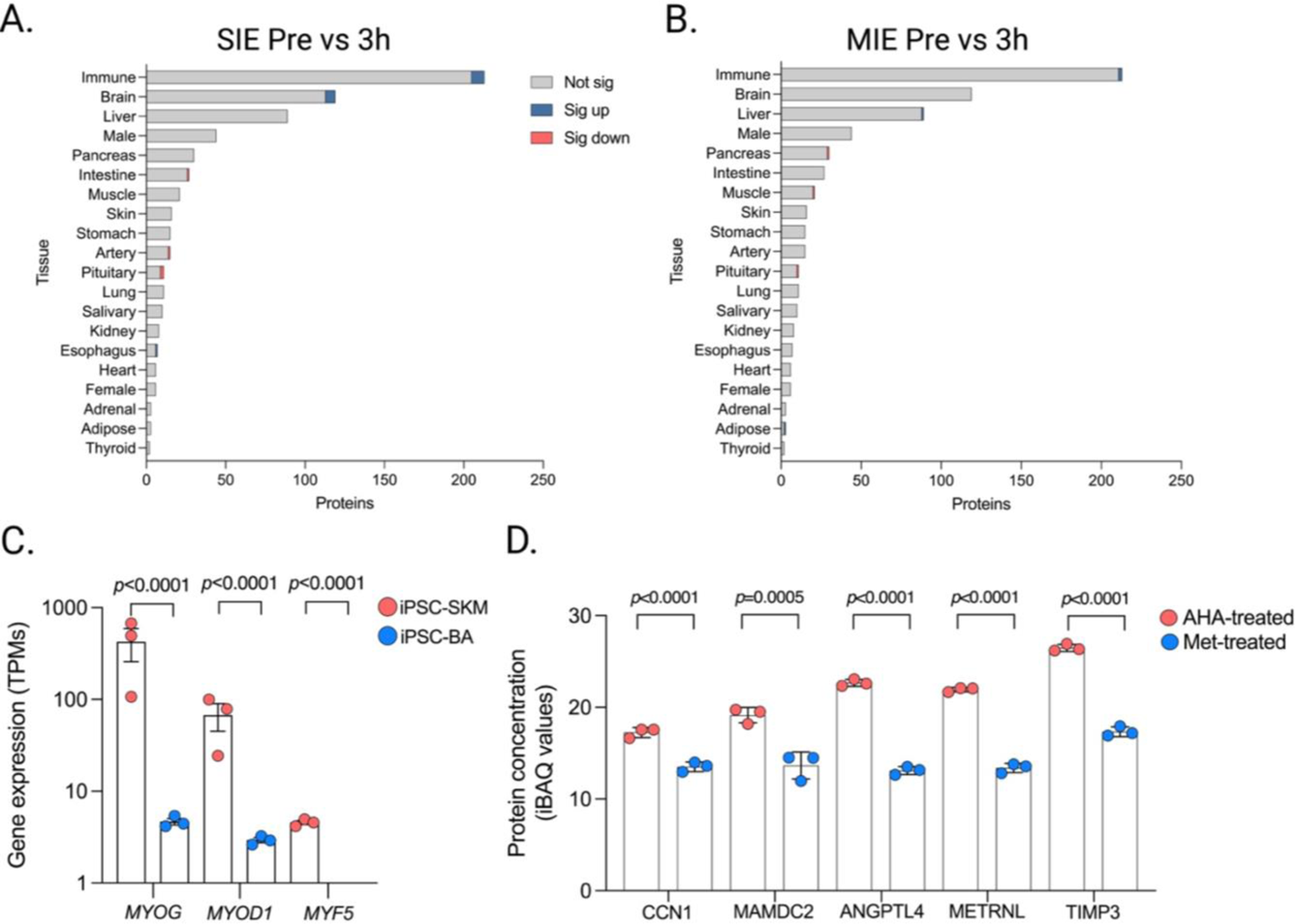
Identifying the organ of origin for exercise regulated plasma proteins and differentially expressed skeletal muscle genes. **A.** All organ-enriched proteins with those increased (blue bar), decreased (red bar), or no change (grey bar) at 3h following SIE. **B.** All organ-enriched proteins with those increased (blue bar), decreased (red bar), or no change (grey bar) at 3h following MIE. **C.** Transcripts per million (TPMs) in iPSC-SKM or iPSC-BA for muscle genes myogenin (*MYOG*), myogenic differentiation 1 (*MYOD1*), and myogenic factor 5 (*MYF5*). **D.** Relative enrichment of proteins based on iBAQ concentrations in AHA- and Met-treated primary human muscle cells. Error bars indicate standard error of the mean. Statistical analysis is described in the methods.

**Figure S5.**
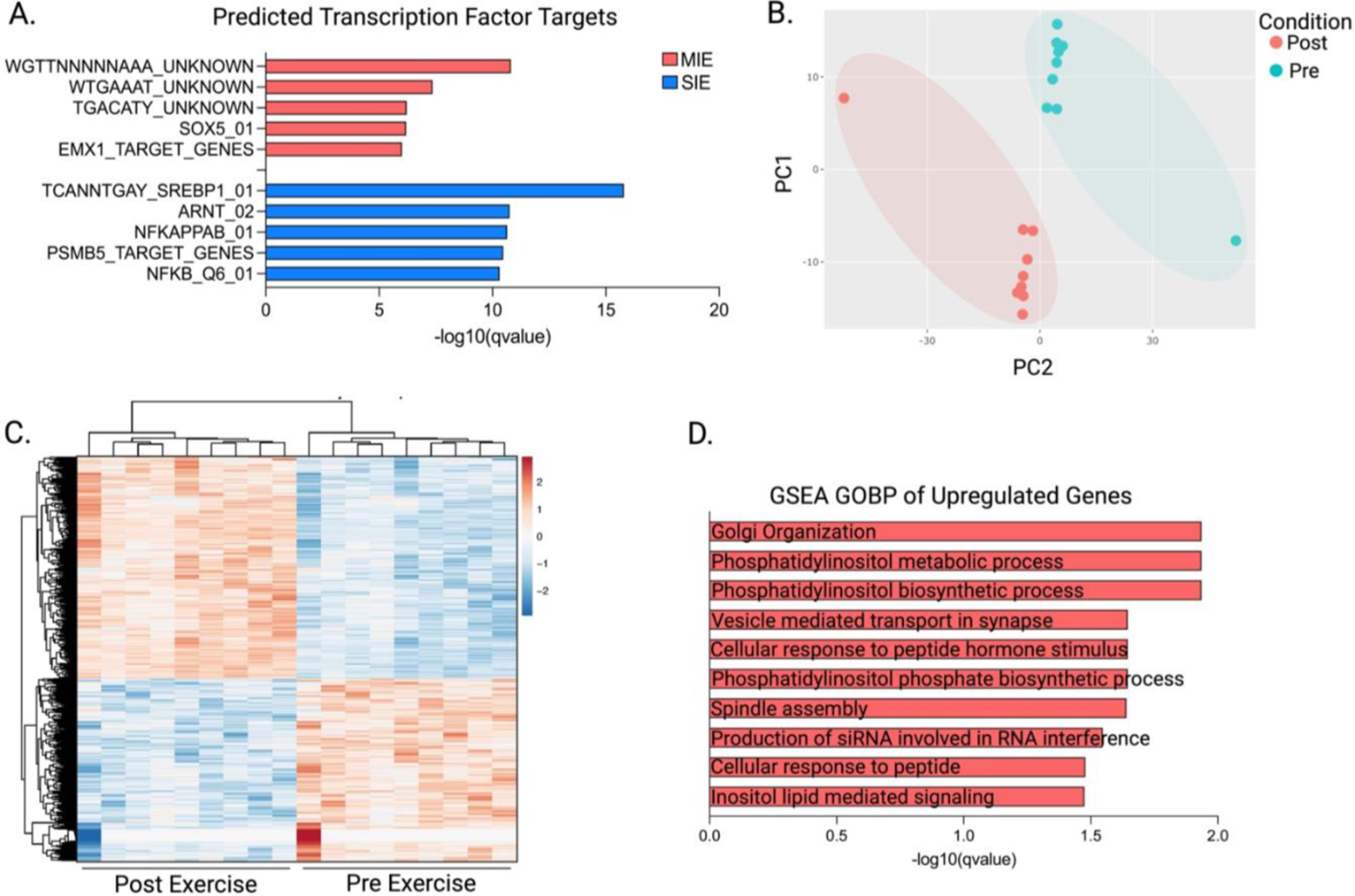
Transcriptomic remodeling of adipocytes and bulk adipose tissue. **A**. Bar graph of the predicted transcription factor targets regulating the differentially expressed adipocyte genes following SIE or MIE plasma. **B**. Principal component analysis of the bulk adipose transcriptome before and after a graded treadmill exercise test. **C**. Heatmap of all differentially expressed bulk adipose genes following a graded treadmill exercise test**. D**. Gene Set Enrichment Analysis (GSEA) of Biological Pathways for all upregulated genes following a graded treadmill exercise test.

